# Pellino 1 Communicates Intercellular Signaling in Chronic Skin Inflammatory Microenvironment

**DOI:** 10.1101/334433

**Authors:** Suhyeon Kim, Seoyoon Bae, Jihyun Park, Geun-Hyoung Ha, Kyungrim Hwang, Hyun-Soo Kim, Jae-Hoon Ji, Heounjeong Go, Chang-Woo Lee

**Affiliations:** Department of Molecular Cell Biology, Samsung Medical Center, Sungkyunkwan University School of Medicine, Suwon 16419, Republic of Korea; Department of Health Sciences and Technology, SAIHST, Sungkyunkwan University, Seoul 06351, Republic of Korea; Genomic Instability Research Center, Ajou University School of Medicine, Suwon 16499, Republic of Korea; Department of Pathology, University of Ulsan College of Medicine, Asan Medical Center, Seoul 05505, Republic of Korea

**Keywords:** Pellino 1, Psoriasis, Keratinocyte, T helper17, Ubiquitination, Interferon regulatory factor 4, Biomarker

## Abstract

Chronic skin inflammation including psoriasis is a multisystem disease, affecting more than 5% of the general population. Here we show that Pellino 1 (Peli1), a signal-responsive ubiquitin E3 ligase, is highly up-regulated in human psoriatic skin lesions and that increased Peli1 expression correlates with the immunopathogenesis of psoriasis-like chronic skin inflammatory disease. Interestingly, Peli1 directly interacts with interferon regulatory factor 4 (IRF4, a transcription factor that plays pivotal roles in proliferation and cytokine production) and induces lysine 63-mediated ubiquitination. Peli1-mediated IRF4 ubiquitination appears to be a common systemic signaling mechanism shared by lesional keratinocytes, dendritic cells, macrophages, and T cells, generating a feedback relationship between keratinocyte and Th17 cell responses. Conversely, inhibition of Peli1 interferes with IRF4 induction and attenuates immunopathogenic signaling in the psoriasis. In summary, Peli1-mediated ubiquitination is a common immunopathogenic intercellular signaling in psoriasis-like chronic skin inflammatory microenvironment. Thus, targeting Peli1 could be used as a potential strategy for psoriasis treatment.

## Introduction

One of the most common immune-mediated chronic inflammatory skin diseases is psoriasis (Lowes *et al*, 2007; Lowes *et al*, 2013). Dysregulation of skin inflammatory signaling resulting in keratinocyte hyperproliferation is one key pathogenic mechanism underlying the development of psoriasis (Harden *et al*, 2015). Keratinocytes are rich sources of antimicrobial peptides, including LL-37, β-defensins, and S100A7 (psoriasin) with chemotactic and immune cell-regulating functions (Buchau *et al*, 2007; Nestle *et al*, 2009). These peptides are also responsive to production of key dendritic cell (DCs)-and T cell-derived cytokines, including interferon-γ (IFNγ), tumor necrosis factor-α (TNFα), IL-17, and IL-20 family of cytokines which in turn generate proinflammatory cytokines (e.g., IL-1, IL-6, and TNFα) and chemokines [e.g., CXC ligand [CXCL]8, CXCL10, and chemokine CC ligand [CCL]20] that contribute to the development of psoriasis (Buchau & Gallo, 2007; Harden et al., 2015; Lowes et al., 2007). However, growing evidence indicates that T cells, particularly those with T helper (Th)17 and Th1 polarization, are heavily present in psoriatic lesions. For instance, interleukin (IL)-23/Th17 axis and Th17-related cytokines play critical roles in the development of psoriasis (Harden et al., 2015; Lowes *et al*, 2014). Therefore, psoriasis arises through chronic interactions between hyperproliferative keratinocytes and infiltrating activated immune cells. Nevertheless, little is known about immunological circuits that maintain established psoriasis lesions.

In psoriatic skin lesions, both αβ and γδ T cells produce IL-17 while activated DCs produce IL-23 and IL-12. They stimulate three populations of resident T cells: Th17, Th22, and Th1 cells (Lowes et al., 2013; Nestle et al., 2009). IL-23 activates Th17 cells to produce IL-17A and IL-17F that drive keratinocyte hyperactivation. It has also been reported that DCs play an essential role in the conversion of nonlesional to lesional psoriatic skin in a transplantation model (Nestle *et al*, 2005). Thus, T cell-and DC-derived cytokines can act on epidermal keratinocytes as proximal inducers of immunogenic circuits in psoriasis. Once activated, skin epidermal cells can produce abundant cytokines, chemokines, and other inflammatory mediators including CXCL8, monocyte chemotactic protein-1/CCL2, CXCL1-3, and CCL20 (Bromley *et al*, 2008; Kennedy-Crispin *et al*, 2012; Zaba *et al*, 2009). However, selective activation of immune cells alone does not seem to be sufficient for initiation of psoriasis. The development of psoriasis requires cross-talk between innate immunity and adaptive immune response.

Interferon regulatory factor 4 (IRF4) is a transcription factor that plays pivotal roles in immune response. Initially, IRF4 was known to be essential for the differentiation of most known CD4^+^ T cell subsets (Huber *et al*, 2014). For example, it facilitates pro-inflammatory cytokine production and Th17 differentiation (Boddicker *et al*, 2015; Brustle *et al*, 2007). The relevance of IRF4 to the development of Th17 cells has been demonstrated in several autoimmune disease models. For example, IRF4 deficient mice are totally resistant to the induction of experimental autoimmune encephalomyelitis (EAE). Such resistance is correlated with the lack of Th17-cell differentiation (Brustle et al., 2007). Conversely, increased IRF4 activity contributes to the development of autoimmune rheumatoid arthritis and large-vessel vasculitis accompanied by elevated synthesis of IL-17 and IL-21 (Chen *et al*, 2008). Consistent with these findings, IRF4 levels are augmented in patients with inflammatory bowel disease and correlated with enhanced production of IL-17 and IL-22 (Mudter *et al*, 2008; Mudter *et al*, 2011). In addition, IRF4 promotes migration of resident DCs from skin to local lymph nodes which triggers T cell-mediated immune responses during cutaneous infection and autoimmune disease (Bajana *et al*, 2012). A recent study on DC differentiation and function has revealed that IRF4-expressing DCs are specialized for instructing IL-17 responses in both mouse and human (Schlitzer *et al*, 2013), indicating that that IRF4 is essential for Th17-mediated autoimmunity. Considering the central function of IRF4 in the differentiation and function of various immune cells, regulation of its expression could be a valuable tool to modulate immune responses.

Pellino family (Peli) proteins are signal-responsive E3 ubiquitin ligases. They are emerging as important players in innate immunity, tumorigenesis, and potentially metabolism (Humphries *et al*, 2015; Park *et al*, 2014). Recent studies have unveiled a critical role of Peli1 in activating toll-like receptor (TLR) and/or T-cell receptor (TCR) signaling-mediated proinflammatory gene expression (Chang *et al*, 2011; Chang *et al*, 2009; Jin *et al*, 2012; Moynagh, 2014). Notably, Peli1 expression is highly suppressed in normal or non-pathological situations. In contrast, pathogenic conditions promote Peli1 expression. For instance, Peli1 expression is upregulated in patients with neutrophilic asthma (Baines *et al*, 2011) and those with diffuse large B cell lymphoma (Park et al., 2014). Moreover, Peli1 expression is associated with poor outcome in B-cell lymphomas (Park et al., 2014), suggesting that constitutive expression of Peli1 contributes to the development of chronic inflammatory immune disease and tumors. Interestingly, Peli1 expression is activated in response to various receptor-mediated signaling events such as those associated with TLR, TCR, and BCR (Chang et al., 2011;Chang et al., 2009; Schauvliege *et al*, 2006). Thus, aberrant regulation of these receptor-mediated signaling pathways can cause unbalanced activation or expression of Peli1 which triggers forced signal cascades, ultimately contributing to the development of certain diseases. In this study, using genetic gain-and loss-of-function approaches, we found that Peli1-mediated ubiquitination was a common immunopathogenic intercellular process. It could be an effective therapeutic biomarker of psoriasis and potentially other chronic skin inflammatory diseases.

## Results

### Peli1 is frequently upregulated in human psoriatic plaques

From our initial approach to examine the expression profile of Peli1 protein in chronic inflammatory skin diseases, Peli1 expression was characterized by comparing non-lesional and lesional skins of psoriasis patients through RNA profiling based on data retrieved from open database analytics (Nair *et al*, 2009). The expression of Peli1 was barely detectable in most healthy human skin samples (Fig 1A). However, it was highly upregulated in lesional skin samples from psoriasis patients. In non-lesional skin samples, expression levels of Peli1 were similar to those in control healthy skin samples (Fig 1A). To further examine the expression profile of Peli1 protein in chronic inflammatory skin diseases, we collected human skin samples from psoriasis patients and heathy controls and compared Peli1 levels. In healthy skin samples, Peli1 was mainly expressed in the cytoplasm of epidermal keratinocytes of the basal layer. It was also expressed in a few dermal endothelial cells and immune cells (Fig 1B). However, in skin plaque lesions from psoriasis patients, Peli1 expression was upregulated in the cytoplasm and a few nuclei of the entire layer of epidermal keratinocytes except stratum corneum (Fig 1B). Peli1 was expressed in infiltrated immune cells with moderate and high intensity. It was also expressed in endothelial cells (Fig 1B). A strong upregulation of Peli1 expression was observed in the entire epidermis, including infiltrated immune cells and blood vessels of skin lesions from psoriasis patients compared to that in healthy skin (Fig 1C), raising the possibility that Peli1 expression is highly correlated with the development of psoriasis.

**Figure 1.**
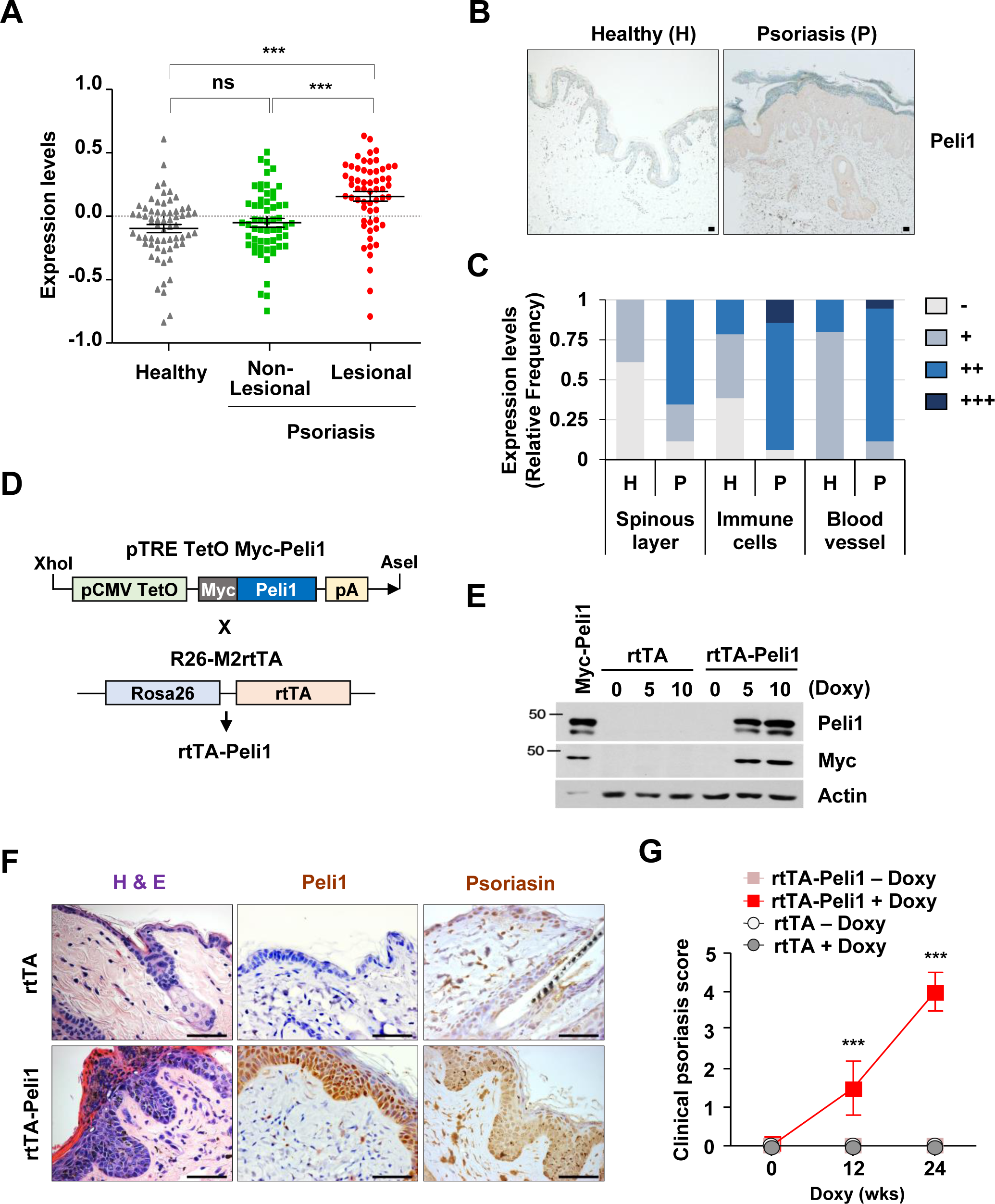
Correlation of Peli1 expression with the development of psoriasis-like skin inflammatory disease. **A.** Comparison of Peli1 mRNA levels in healthy human skin (n = 64), non-lesional skin (n = 58), and lesional skin (n = 58) samples of psoriasis patients. Microarray data sets processed with Robust Multichip Average (RMA) method were retrieved from Gene Expression Omnibus (GEO) Database (reference 30, accession number: GSE 13355). Expression values in dot graph are adjusted RMA expression values (log scale) to account for batch and sex effects. Data are presented as means ± SEM. * *P* < 0.05, ** *P*< 0.01, *** *P* < 0.001; ns: not significant based on Student’s *t*-test. **B.** Normal human skin (breast) immuno-stained for Peli1 (left 40x). Psoriasis lesion immuno-stained for Peli1 (right, 40x). Scale bars, 50 μm. **C.** Peli1 expression was evaluated semi-quantitatively by comparing expression intensity between healthy skin (H) and psoriatic skin lesions (P) in spinous layer and infiltrated immune cells as described in methods. **D.** Generation of doxycycline-inducible human Peli1 transgenic mice. cDNA sequence for human Peli1 was placed under Tet-responsive promoter and introduced into fertilized mouse oocytes. To generate rtTA-Peli1 mice, founders were crossed with R26-M2rtTA mice (B6.Cg-Gt(ROSA)26Sor^™1^(rtTA*M^2^)Jae/J). For inducibility, Myc epitope-tagged human Peli1 gene sequences under the control of TetO promoter and human early cytomegalovirus enhancer were included. **E.** Immunoblotting of tissues from rtTA and rtTA-Peli1 mice using Peli1 and Myc antibodies revealing Peli1 protein expression. Doxy: doxycycline trea™ent. **F.** Adult rtTA control mice and rtTA-Peli1 inducible transgenic mice were treated with or without doxycycline (2 mg/ml) for 12 weeks.H&E-staining and immunostaining of representative skin sections for Peli1 and Psoriasin. Scale bars, 50 μm. **G.** Psoriasis incidence in rtTA and rtTA-Peli1 mice at 12 weeks after doxycycline trea™ent. rtTA-doxy: rtTA without doxycycline trea™ent (n = 30); rtTA-Peli1-doxy: rtTA-Peli1 without doxycycline trea™ent (n = 35); rtTA+doxy: rtTA with doxycycline trea™ent (n = 62); rtTA-Peli1+doxy: rtTA-Peli1 with doxycycline trea™ent (n = 78). Clinical psoriasis scores of rtTA and rtTA-Peli1 mice treated with or without doxycycline are shown. Scores were obtained from 3 to 5 mice per group. Data are presented as means ± SEM. *** *P* < 0.001, two-way ANOVA.

### Activation of Peli1 expression causes epidermal hyperplasia and psoriasis-like skin inflammation

Since Peli1 was frequently upregulated in psoriatic skin lesions, we generated double positive mice carrying both human Peli1 transgene and rtTA inducer allele (Peli1^+^/ROSA26-rtTA^+^, referred to herein as doxycycline inducible Peli1 transgenic mice, hereafter rtTA-Peli1) (Fig 1D). Immunoblotting showed detectable levels of human Peli1 protein only in double-transgenic rtTA-Peli1 by doxycycline trea™ent (Fig 1E). However, non-transgenic rtTA mice treated with doxycycline did not express any human Peli1 (Appendix Fig S1A). Doxycycline-inducible rtTA-Peli1 mice showed no detectable level of human Peli1 protein when they were left untreated (Fig 1E). Offspring of three different rtTA-Peli1 founder lines exhibited the highly similar macroscopic pathologies as early as 10-12 weeks after doxycycline administration (Appendix Fig S2A, B). By 12 weeks post doxycycline trea™ent, more than 95% of rtTA-Peli1 mice showed many psoriasis-like features, including extensive erythema, hair loss, sever pruritus, and loosely adherent silver-white scaling (Appendix Fig S2A, B). Histological analysis of skin samples from treated 3-month-old rtTA-Peli1 mice revealed acanthosis, regularly elongated rete ridges, thickened cornified layer (hyperkeratosis), epidermal hyperplasia (acanthosis), and parakeratosis (Fig 1F). The severity of symptoms in the rtTA-Peli1 mice based on human psoriasis-like area and severity index (PASI) was correlated with the duration of Peli1 overexpression (Fig 1G). However, rtTA mice treated with doxycycline and rtTA-Peli1 mice without doxycycline did not show any sign of skin inflammation (Fig 1G). Further immunohistochemistry showed an increase in numbers of keratinocytes expressing K14 (expressed in mitotically active basal epidermal layer cells), K10 (expressed in abnormally differentiated cells in epidermal layer), loricrin (a terminally differentiating structural protein comprising of cornified envelope), and Ki67 (strictly associated with cell proliferation) in mice overexpressing human Peli1 protein (Fig 2A). In addition, increased epidermal infiltrations of CD3^+^ T cells and F4/80^+^ macrophage and expansion of CD31^+^ endothelial cells were observed in rtTA-Peli1 mice compared to those in rtTA mice treated with doxycycline (Fig 2A, B). Results of immunohistochemical analysis of rtTA-Peli1 showed high similarity with phenotypes of human psoriatic skin (Table 1) (Boehncke, 2015; Sano *et al*, 2005).

**Figure 2.**
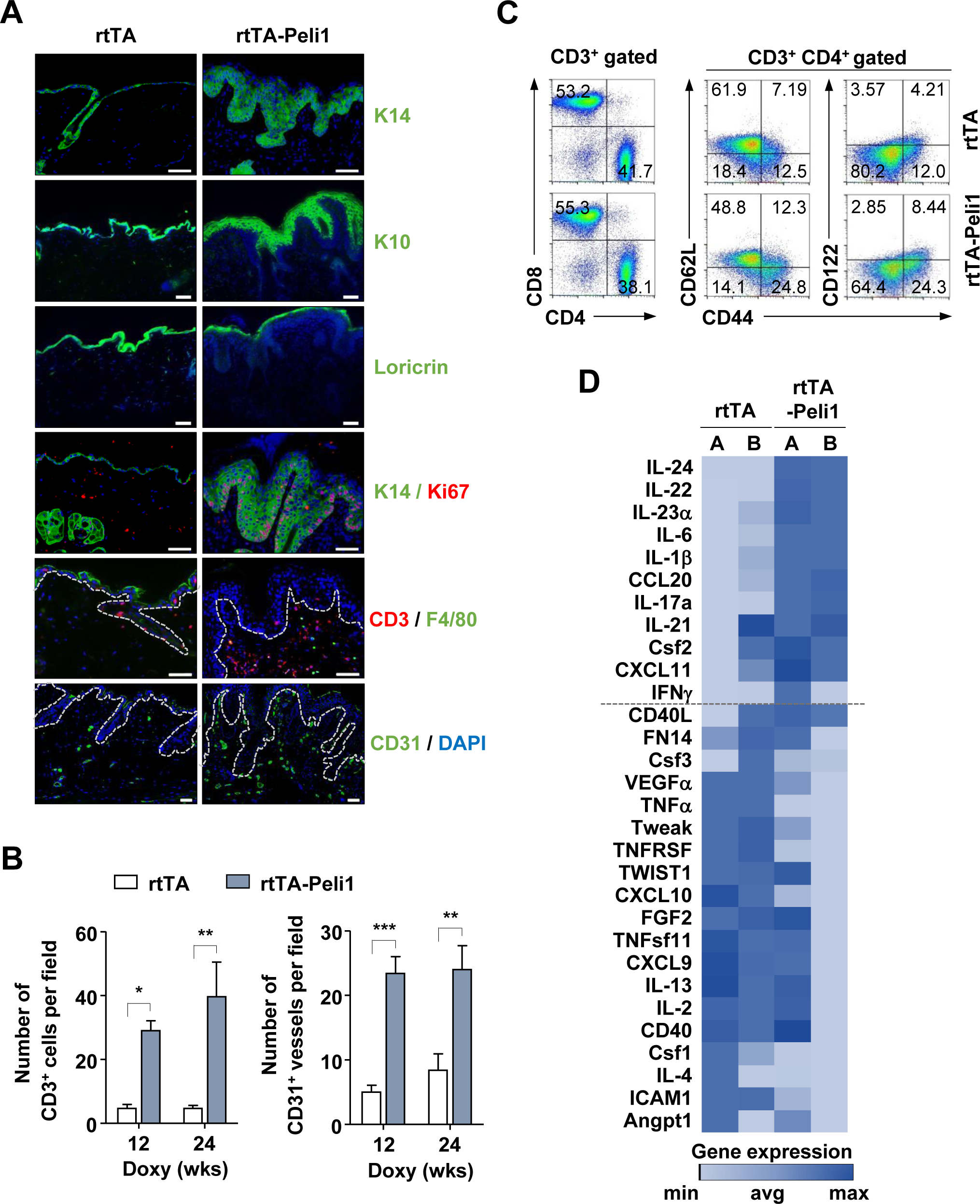
Development of epidermal hyperplasia and psoriasis-like skin inflammation in inducible transgenic mice expressing human Peli1 gene. **A.** Adult rtTA control mice and rtTA-Peli1 inducible transgenic mice were treated with doxycycline (2 mg/ml) for 12 weeks. Representative immunofluorescence images of skin sections for keratinocyte differentiation markers (keratin 14, keratin 10, and loricrin), proliferation marker (Ki67), dermal infiltrated immune cells markers (CD3 and F4/80), and angiogenesis marker (CD31) are shown. Dotted lines indicate the border between epidermis and dermis. Scale bars, 50 μm. **B.** Quantification of CD3^+^ cells and CD31^+^ vessels based on five skin sections from five independent mice with indicated genotypes (n = 3 mice per genotype). **C.** Flow cytometry of T cells from rtTA and rtTA-Peli1 mice following trea™ent with doxycycline for 24 weeks. Immunocytes were isolated from draining lymph-nodes (inguinal, axillary, brachial, and cervical lymph-nodes) of rtTA and rtTA-Peli1 mice. Cells were stained with CD3, CD4, CD44, CD62L, and CD122 followed by flow cytometry analysis. Gated CD3^+^CD4^+^ T cells were analyzed for CD62L, CD44, and CD122 expression. **D.** Heat map depicting real-time quantitative PCR (qRT-PCR) analysis of inflammation-related genes in skin samples from doxycycline-treated rtTA and rtTA-Peli1 mice (two mice for each genotype). Genes were ranked based on fold-change in expression. Genes above the dashed line were highly elevated in lesional skin samples from mice overexpressing Peli1 (rtTA-Peli1 with doxycycline trea™ent).

**Table 1.**
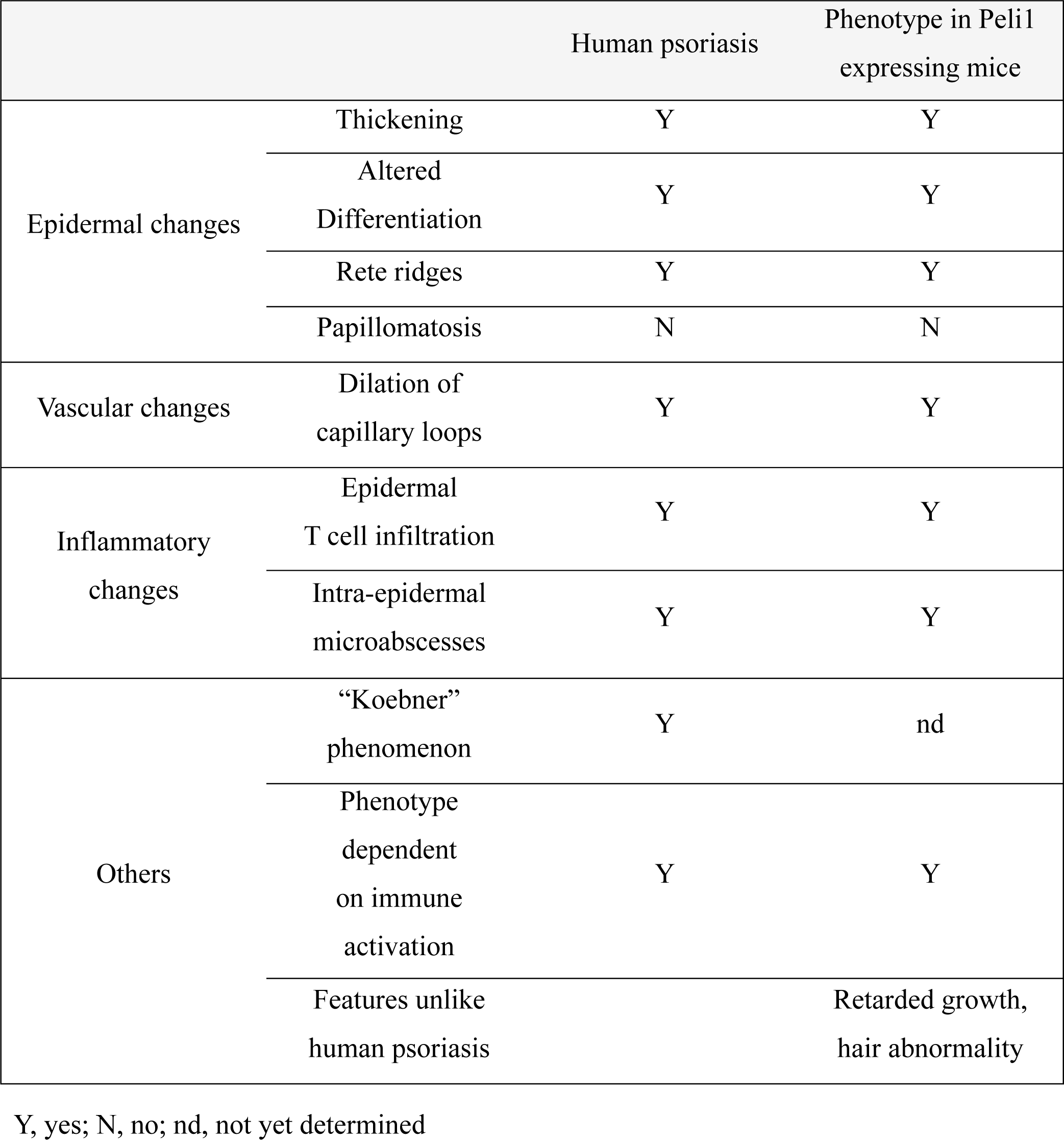
Comparison of pathological phenotype similarity between human psoriasis and Peli1 expressing mice

Psoriatic lesions are characterized by the infiltration of T cells that secrete cytokines such as IFNγ, TNFα, IL-17, and IL-23(Lowes et al., 2007). Since skin draining lymph-nodes of doxycycline treated rtTA-Peli1 mice were enlarged with more cellularity (Appendix Fig S3A, B), we assessed the activation status of CD4^+^ T cells based on surface expression levels of their activation markers CD44, CD62L, and CD122. Untreated rtTA-Peli1 mice did not show obvious abnormalities in the frequency of naïve or activated/memory T cells (Appendix Fig S3C). However, doxycycline treated rtTA-Peli1 mice showed higher frequency of activated/memory T cells with fewer naïve T cells in skin draining lymph-nodes (Fig 2C and Appendix Fig S3D). CD8^+^ T cells also exhibited a more activated/memory phenotype in rtTA-Peli1 mice compared to those in rtTA mice. Consistent with this phenotype, CD4^+^ T or CD8^+^ T cells in 24-week-old Peli1 inducible Tg mice had more activated/memory T cells in both lymph nodes and spleen (data not shown). Furthermore, more IL-17^+^ and IL-22^+^CD4^+^ T cells in skin draining lymph-nodes of doxycycline treated rtTA-Peli1 mice than those in their counterparts were observed (Appendix Fig S3E).

It is known that psoriatic plaques are due to dysregulated interactions of innate and adaptive components of the immune system affecting resident immune cells of the skin(Boehncke, 2015). To gain insight into the molecular pathogenesis of Peli1-induced psoriasis, we analyzed a panel of key inflammation-related genes usually activated during psoriasis using an RT^2^ profiler PCR array (Fig 2D). We found significant changes in expression levels of several Th17-related cytokines such as cytokines secreted by Th17 cells (IL-17a, IL-21, IL-22, and IL-24), cytokines involved in Th17 differentiation (IL-21 and IL-23), and chemokine CCL20 for trafficking of Th17 cells. Skin samples of rtTA-Peli1 mice showed strong up-regulation of IL-17a (6.5-fold), IL-21 (1.7-fold), IL-22 (12.5-fold), IL-24 (10.9-fold), IL-23(1.8-fold), and CCL20 (1.7-fold) compared to those of rtTA mice (Fig 2D). Taken together, our results strongly suggest that increased Peli1 expression initiates the development of chronic skin inflammatory disease, similar to phenotype of human psoriatic skin.

### Peli1 mediates the development of psoriasis-like disease by reciprocal cross-linking between keratinocyte and Th17 CD4^+^ T cell response

To determine whether psoriasis developed in rtTA-Peli1 mice could be ameliorated by Peli1 inhibition, mice were withdrawn from doxycycline trea™ent (Fig 3A). rtTA and rtTA-Peli1 mice were administrated with doxycycline for 12 weeks, after which one group continued to receive doxycycline for an additional 10 weeks (ON) while the second group was switched to water (OFF). Four weeks after doxycycline withdrawal,expression of Peli1 was barely detectable in most organs tested (data not shown). Withdrawal of doxycycline (ON → OFF) in rtTA-Peli1 mice led to sharp recovery from psoriasis. However, continuous trea™ent with doxycycline (ON → ON) worsened the psoriasis based on PASI scoring (Fig 3B) and H&E staining (Fig 3C, upper panels). Immunohistochemistry staining further revealed that removal of doxycycline trea™ent to rtTA-Peli1 mice dramatically decreased numbers of keratinocytes expressing K14 and PCNA (Fig 3C, lower panels), indicating that inhibition of Peli1 reversed psoriasis in these mice. Furthermore, the removal of doxycycline trea™ent to rtTA-Peli1 mice resulted in significant reductions in expression levels of Th17-related inflammatory genes, including IL-17a, IL-21, IL-22, IL-24, IL-23, and CCL20 (Fig 3D). Taken together, these data indicate that Peli1 is critically involved in the development of psoriasis by mediating the cross-link between keratinocyte and Th17 CD4^+^ T cell response.

**Figure 3.**
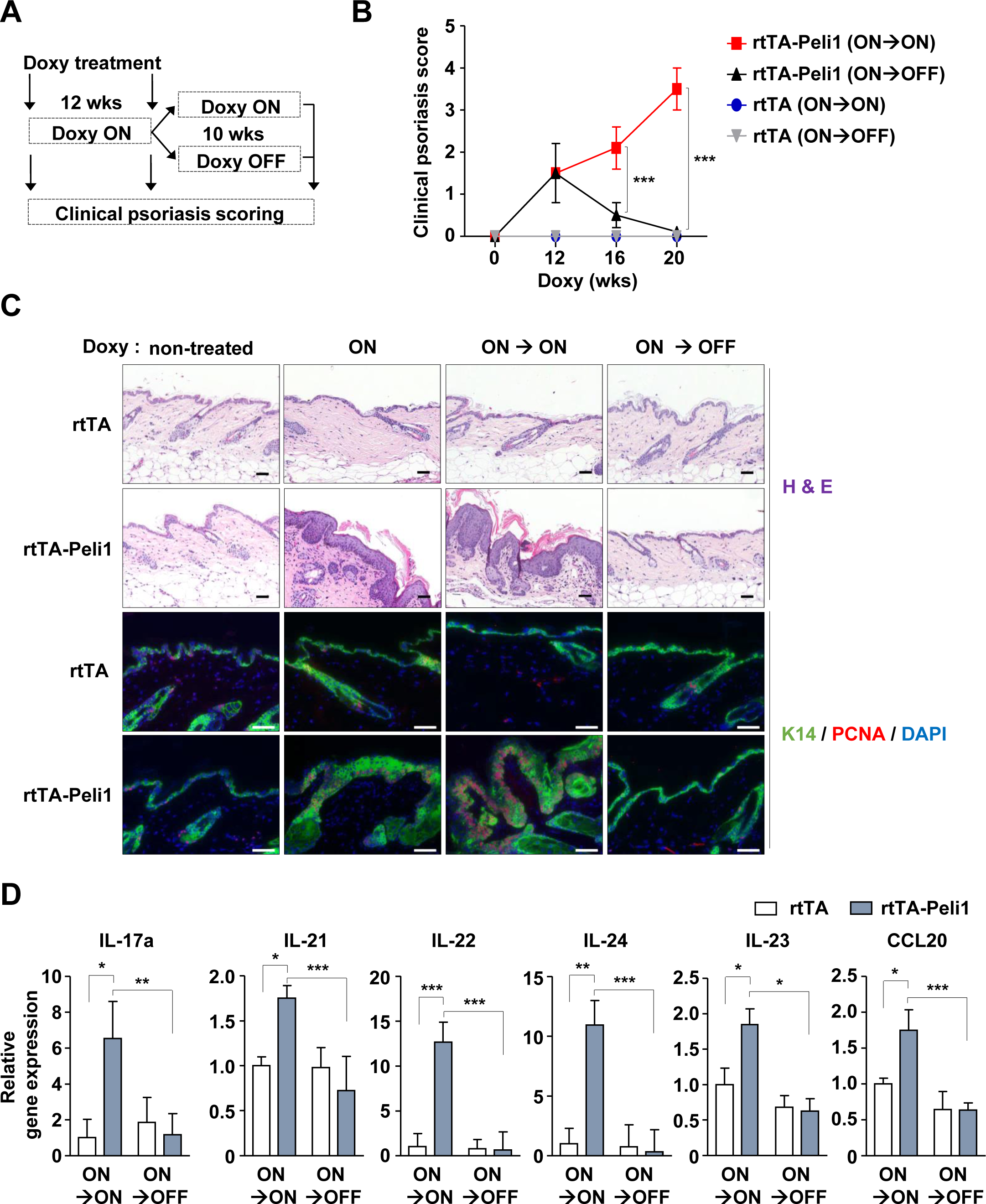
Peli1 is involved in the development of psoriasis-like disease and Th17 cell response. **A.** Schematic illustration of the experimental design using doxycycline-inducible transgenic mice with different Doxy administration conditions. rtTA and rtTA-Peli1 mice were administrated with doxycycline for 12 weeks. Each group of doxycycline-administered mice were then further treated with (ON) or without (OFF) doxycycline for an additional 10 weeks. Doxy: doxycycline. **B.** Comparison of clinical psoriasis scores of doxycycline administered rtTA and rtTA-Peli1 mice after continued trea™ent (ON→ON) or removal (ON→OFF) of doxycycline at indicated time points. Scores were from 3 to 5 mice per group. Data are presented as means ± SEM. *** *P* <0.001, two-way ANOVA. **C.** Hematoxylin and eosin (top) and immunohistochemical staining of representative skin sections for keratin 14 and PCNA expression (bottom). Scale bars, 50 μm. **D.** Comparison of mRNA expression levels of pro-inflammatory cytokines. Quantitative RT-PCR for cytokine expression was performed using skin tissues. Data are presented as means ± SEM (n = 3). * *P* < 0.05, ** *P* < 0.01, *** *P* <0.001; ns: not significant, two-way ANOVA.

### Overexpression of Peli1 in epidermal cells initiates the development of psoriasis-like disease

To dissect pathways by which Peli1 expression led to the development of psoriasis, we utilized the adoptive transfer function of congenic mouse bone marrow (BM) cells. BM cells were isolated from donor mice and injected intravenously into lethally irradiated recipient rtTA and rtTA-Peli1 mice (Fig 4A) or Pepboy.1 mice (Fig 4B). After BM transfer, each group was analyzed for psoriasis-like phenotypes at 12 weeks post administration of doxycycline. Interestingly, only rtTA-Peli1 chimera mice receiving Pepboy.1 BM cells (Group 2) began to develop psoriatic-like lesions, including epidermal thickening, scaling, small rete ridge, and abnormal proliferative basal keratinocytes of the epidermis (Fig 4C). We also found significant abnormality in the ratio of naïve to activated T cells in rtTA-Peli1 mice receiving pepboy.1 BM cells (Group2) (Fig 4D). To determine percentages of Th1, Th2, and Th17 cytokine-positive cells in skin draining lymph-nodes, we also analyzed expression levels of intracellular IFNγ, IL-4, IL-17a, and IL-22 by flow cytometry. Of interest, rtTA-Peli1 mice receiving pepboy.1 BM cells (Group2) showed increase in percentage of IL-17^+^IL-22^+^CD4^+^ T cells compared to mice from other experimental groups (Fig 4D).

**Figure 4.**
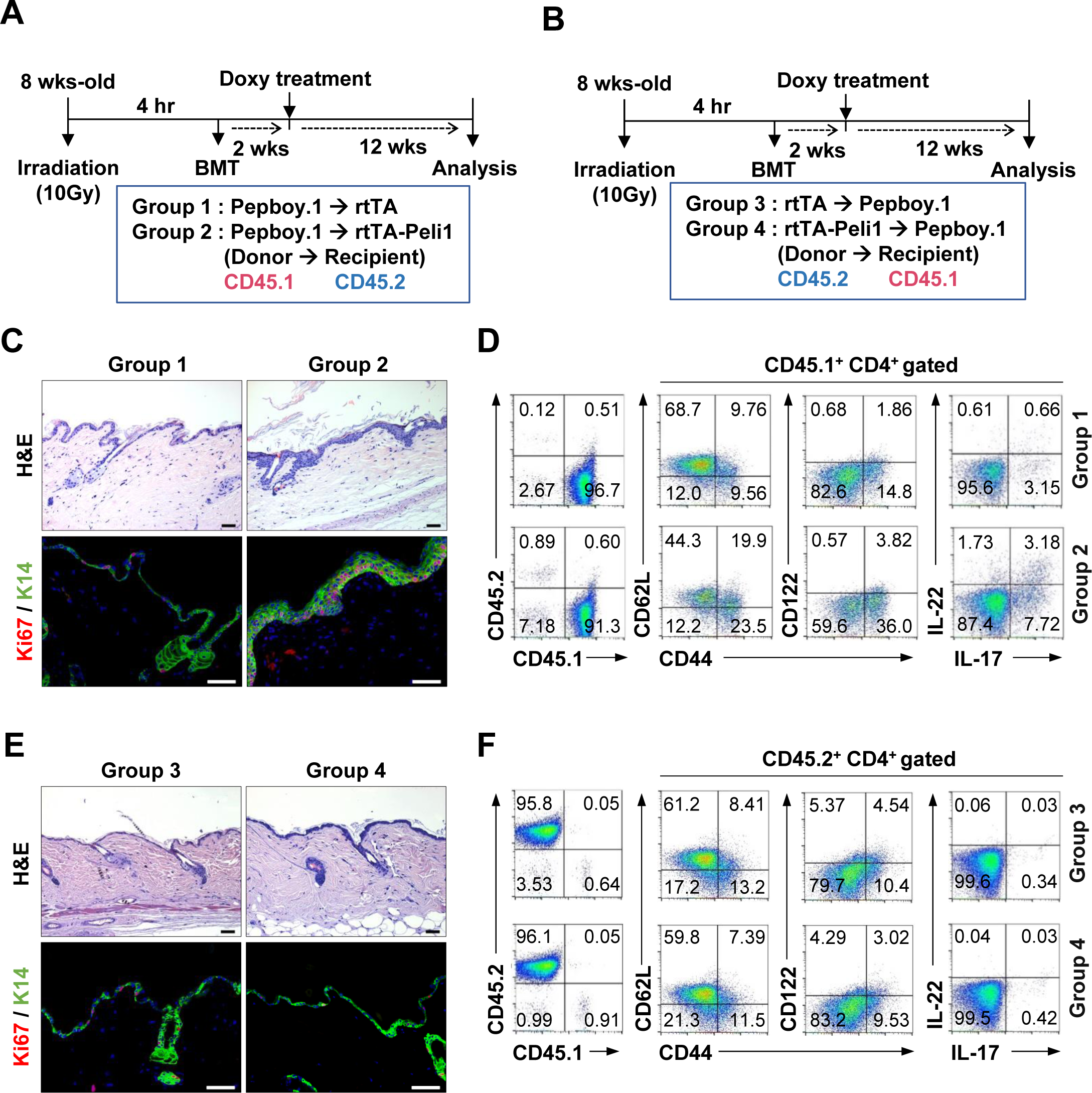
Overexpression of Peli1 in epidermal cells but not in T cells triggers the development of psoriasis-like disease. **A, B.** Schematic representation showing the generation of chimeric recipient mice. Bone marrow (BM) cells were isolated from donor mice and injected into lethally irradiated recipient mice. The following four groups were generated: Pepboy.1 (CD45.1) BM →rtTA (CD45.2) mice (Group 1); Pepboy.1 (CD45.1) BM → rtTA-Peli1 (CD45.2) mice (Group 2); rtTA (CD45.2) BM → Pepboy.1 (CD45.1) mice (Group 3); and rtTA-Peli1 (CD45.2) BM →Pepboy.1 (CD45.1) mice (Group 4). After BMT, each group was given doxycycline for 12 weeks and then sacrificed (n = 3 for each group). **C, E.** Histology (top) and immunofluorescence of keratin 14 and Ki67 expression (bottom). Scale bars, 50 μm. **D, F.** Representative FACS plots showing the efficiency of bone marrow transplantation, activation of T cell, and intracellular expression of IL-17 and IL-22. Gating blood cells for CD45.1 and CD45.2 expression revealed more than 90% engraf™ent of donor cells in recipient mice. Lymphocytes were isolated from draining lymph-nodes and stained for CD45.1, CD45.2, CD4, CD44, CD62L, CD122, IL-17, and IL-22 followed by flow cytometry analysis.

Further histopathological analysis showed that pepboy.1 chimeric mice receiving rtTA-Peli1 BM cells (Group 4) had no significant signs of psoriasis-like phenotype (Fig 4E, F), implying that overexpression of Peli1 in lymphocytes only seemed to be insufficient to develop psoriatic lesions. To assess the effect of Peli1 overexpression on T cell activation, we stimulated rtTA and rtTA-Peli1 CD4^+^ T cells with anti-CD3 and anti-CD28 *in vitro*. Our results showed that hyperactivation of T cells was particularly evident in doxycycline-treated rtTA-Peli1 mice whereas overexpression of Peli1 in T cells had no significant effect on the activation of TCR signaling after TCR stimulation (Appendix Fig S4A). To further investigate the differentiation capacity of Peli1-overexpressing T cells, we performed *in vitro* polarization using MACS-purified CD4^+^ T cells activated under Th1, Th2, and Th17 polarization conditions. Our results showed that polarization potentials of Peli1-overexpressing T cells toward their respective Th1 (driven by IL12 and anti-IL4), Th2 (driven by IL-4 and anti-IFNγ), and Th17 (driven by TGFβ, IL-6, anti-IFNγ, and anti-IL4) lineages were similar to those of control T cells (Appendix Fig S4B). Taken together, these data indicate that Peli1-mediated hyperactivation of keratinocytes can initiate pathogenic hyperactivation of T cells in the development of skin inflammation and psoriasis lesion.

### Peli1-mediated IRF4 induction relays intercellular signaling from lesional keratinocytes to infiltrated immune cells

It is known that epidermal keratinocytes are key players in innate immunity that can induce and switch classes of T cells recruited to lesional skin (Buchau & Gallo, 2007; Harden et al., 2015; Nestle et al., 2009). Therefore, we studied the underlying molecular mechanism involved in the activation of keratinocyte proliferation and the development of psoriasis induced by overexpression of Peli1. We first compared cell cycle profiles between rtTA and rtTA-Peli1 epidermal keratinocytes by flow cytometry analysis (Appendix Fig S5A). We also monitored expression of PCNA (a S phase marker), PCNA and phospho-H3^S10^ double expression (a G2 phase marker), and expression of phospho-H3^S10^ (a marker of mitosis) (Appendix Fig S5B, C). Overall cycle progression in rtTA-Peli1 epidermal keratinocytes was significantly accelerated compared to that in same-aged rtTA epidermal keratinocytes, showing significant differences in G0/G1 phase and rapid progression into S, G2, and M phases induced by Peli1 expression.

Next, we determined expression profiles of a series of representative marker molecules involved in signaling pathways of hyperproliferation and hyperactivation of keratinocyte using skin tissues isolated from rtTA and rtTA-Peli1 mice (Fig 5A, B). Healthy skin and Peli1-induced psoriatic skin lesions showed significant differences in several cell cycle progression regulators, including elevated expression of PCNA,Mcm6, Cyclines A, B, D and E, phosphorylation of Rb at serine 807/811 residue, and Aurora B. However, reduction of Cdk inhibitor p27 was observed (Fig 5A), indicating that Peli1 expression could lead to hyperproliferation of lesional keratinocytes. We next compared expression profiles of genes involved in activation of psoriasis-related cytokine signaling pathways such as STAT3, NF-kB, and TLR signaling (Gudjonsson *et al*, 2007). As shown in Fig 5B, STAT3 phosphorylation, cIAP2, and levels of its downstream target proteins TRAF6 and RIP1 were increased in Peli1-induced psoriatic skin lesions, suggesting that Peli1 could activate STAT3 and NF-kB signaling that contribute to the development of psoriasis. However, most genes with altered expression levels in Peli1-induced psoriatic lesion did not show any changes in non-psoriatic lesions of rtTA-Peli1 mice at 7 weeks after doxycycline trea™ent (Fig 5C, D).

Surprisingly, there was a major difference in the induction of IRF4 (a transcription factor expressed in hematopoietic cells that plays a pivotal role in both proliferation and cytokine production (Huber & Lohoff, 2014) between healthy skin and Peli1-induced psoriatic skin (Fig 5B). IRF4 induction was distinct even in non-psoriatic lesion of mice expressing Peli1 (Fig 5D). Recent studies have suggested that IRF4 plays a key role in Th17 differentiation (Alam *et al*, 2014; Brustle et al., 2007; Korn *et al*, 2009). IRF4 can bind to IL-17A promoter and induce its activity (Mudter et al., 2011). Recent studies have also shown that IRF4-expressiong DCs are specialized in driving IL-17 responses (Persson *et al*, 2013; Schlitzer et al., 2013). Furthermore, IRF4 is upregulated in skin lesions of psoriasis patients (Ni *et al*, 2012). A reduction in IRF4 during phototherapy of psoriasis provides beneficial effects (Alam et al., 2014), thereby indicating its crucial role in mediating psoriasis. To investigate how Peli1 might affect IRF4, we compared IRF4 expression in rtTA-Peli1. Continued trea™ent with doxycycline (ON → ON) in rtTA-Peli1 mice increased IRF4 expression in epidermal keratinocytes and populations of IRF4-expressing dermal infiltrating cells. However, withdrawal of doxycycline (ON → OFF) in rtTA-Peli1 mice clearly decreased IRF4 expression (Fig 5E). Next, subsets of dermal infiltrating immune cells involved in the induction of IRF4 by Peli1 expression were determined. Single-cell suspensions from epidermis, dermis, and draining lymph-nodes were prepared and expression levels of IRF4 protein in T cells, macrophages, and dendritic cells were compared (Fig 5F-H). Interestingly, IRF4 expression levels were markedly upregulated in DCs, macrophages, and T cells of rtTA-Peli1 dermal and epidermal lesions, but not in rtTA dermis or epidermis (Fig 5F, G). There were no significant changes in IRF4 expression levels in the same cells from draining lymph-nodes between rtTA and rtTA-Peli1 mice (Fig 5H). Together, these data indicate that Peli1 expression can sequentially activate IRF4 induction from keratinocytes to infiltrated immune cells in the psoriatic microenvironment.

**Figure 5.**
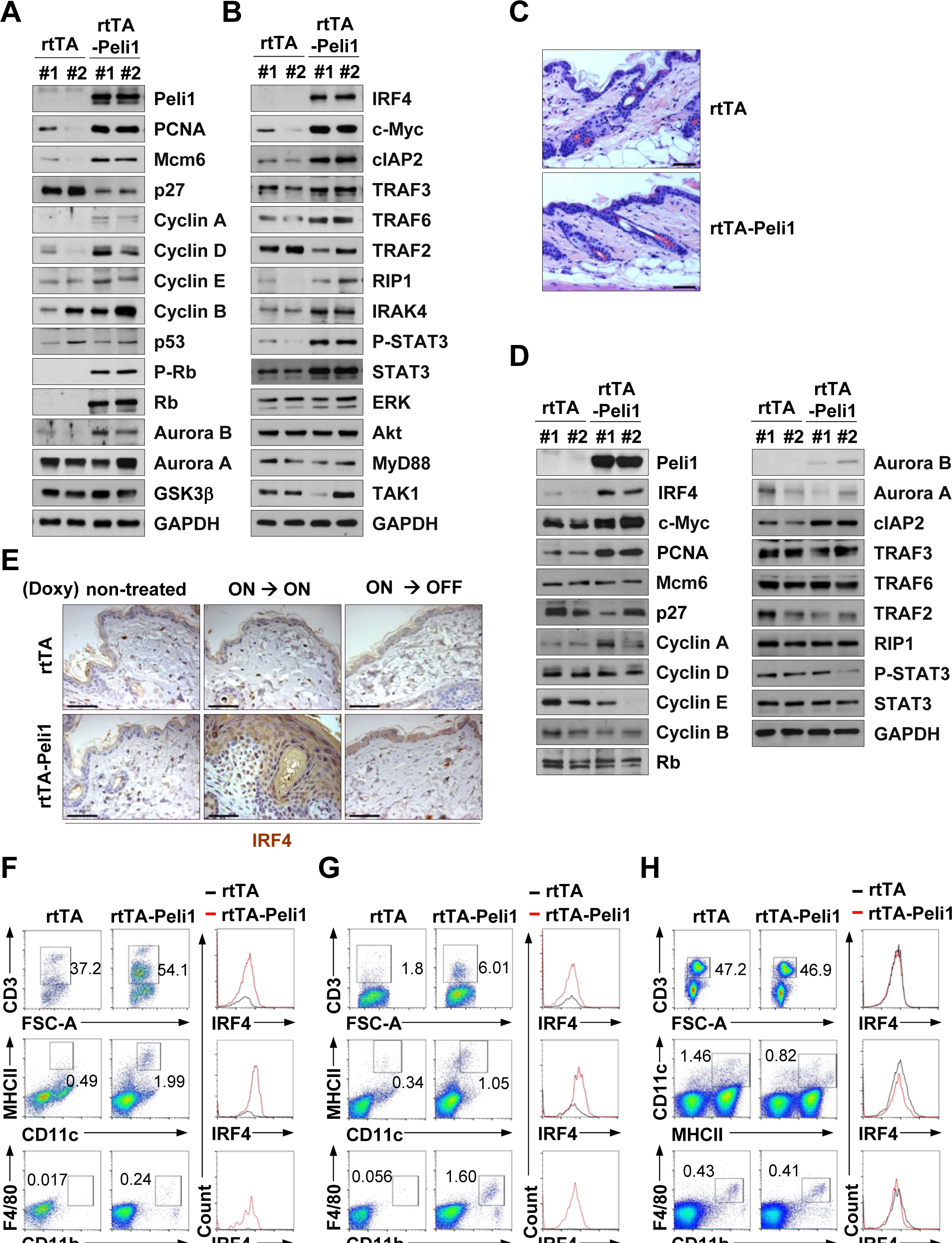
IRF4 is upregulated in Peli1-induced psoriatic intercellular signaling circuits. **A, B.** Immunoblot analysis of indicated protein in skin tissues from rtTA and rtTA-Peli1 mice after 24 weeks of doxycycline trea™ent. **C.** Comparison of protein expression profile between rtTA and rtTA-Peli1 mice during early stage of psoriasis development.Hematoxylin and eosin-stained sections of skin tissues from rtTA and rtTA-Peli1 mice at 7 weeks post doxycycline administration are shown. **D.** Immunoblot analysis of indicated protein in skin tissues from rtTA and rtTA-Peli1 mice at 7 weeks after doxycycline trea™ent. Scale bars, 50 µm. **E.** Immunostaining of skin sections from rtTA and rtTA-Peli1 mice after 24 weeks of doxcycline trea™ent with an IRF4 antibody for the detection of IRF4 induction. Scale bars, 50 μm. **F-H.** Representative FACS plots of single-cell suspensions of epidermal (**F**), dermal (**G**), and draining lymph-nodes (**H**) from rtTA and rtTA-Peli1 mice after 24 weeks of doxycycline trea™ent. Cells were stained with antibodies specific for CD3, CD11c, MHCII, CD11b, F4/80, and IRF4. Histograms show expression levels of IRF4 on T cells (CD3^+^, top panel), DCs (CD11c^+^/MHCII^+^, middle panel), and macrophages (CD11b^+^/F4/80^+^, bottom panel). The percentage of each population is shown in plots.

### Peli1 deficiency delays psoriasis development potentially by inability to induce IRF4

It is well known that topical application of imiquimod (IMQ), an effective agonist for TLR7 in mice (TLR7 and TLR8 in human), can induce and exacerbate psoriasis by activating IL-23/Th17 pathway via TLR7 (Ueyama *et al*, 2014; van der Fits *et al*, 2009). We thus determined whether depletion of Peli1 might counteract the development of psoriasis induced by IMQ trea™ent. To test this possibility, we generated Peli1 mutant mice by a conventional targeting strategy in which coding exon 2 was eliminated by Cre recombinase (Appendix Fig S1B, C). Similar to a previous report (Chang et al., 2011), these newly generated homozygous Peli1 knockout (KO) mice did not show apparent abnormalities in growth or survival (data not shown). We investigated the effect of Peli1 deficiency on the development of IMQ-induced psoriasis and found that Peli1 KO mice displayed significant reductions in IMQ-induced thickening, scaling, and erythema of skin lesional areas (Fig 6A-C). At day 5 after IMQ application, signs of improvement began to occur largely in Peli1 KO mice compared to WT mice (Fig 6B, C). Immunohistochemistry staining demonstrated that deletion of Peli1 in mice treated with IMQ significantly attenuated the development of psoriasis and reduced populations of CD3^+^ T cells, F4/80^+^ macrophages, and cells expressing Ki67 and K14 (Fig 6D, E). This IMQ-induced psoriasis model revealed that levels of IRF4 were significantly decreased in epidermal keratinocytes and dermal infiltrating immune cells of Peli1 KO mice compared to those of Peli1 WT mice (Fig 6F). Furthermore, we compared expression levels of IRF4 protein in T cells, macrophages, and dendritic cells using single-cell suspensions isolated from epidermis, dermis, and draining lymph-nodes between IMQ-treated Peli1 WT and KO mice (Fig 6G-I). Interestingly, Peli1 deficiency significantly downregulated or failed to induce IRF4 levels in DCs, macrophages, and T cells compared to Peli1 WT. There were no significant changes in IRF4 expression levels in cells obtained from draining lymph-nodes between Peli1 WT and KO mice (Fig 6I), similar to data observed in Fig 5H. Taken together, these data indicate that inhibition of Peli1 attenuates the development and disease severity of IMQ-induced psoriasis most likely by interfering with IRF4 induction.

**Figure 6.**
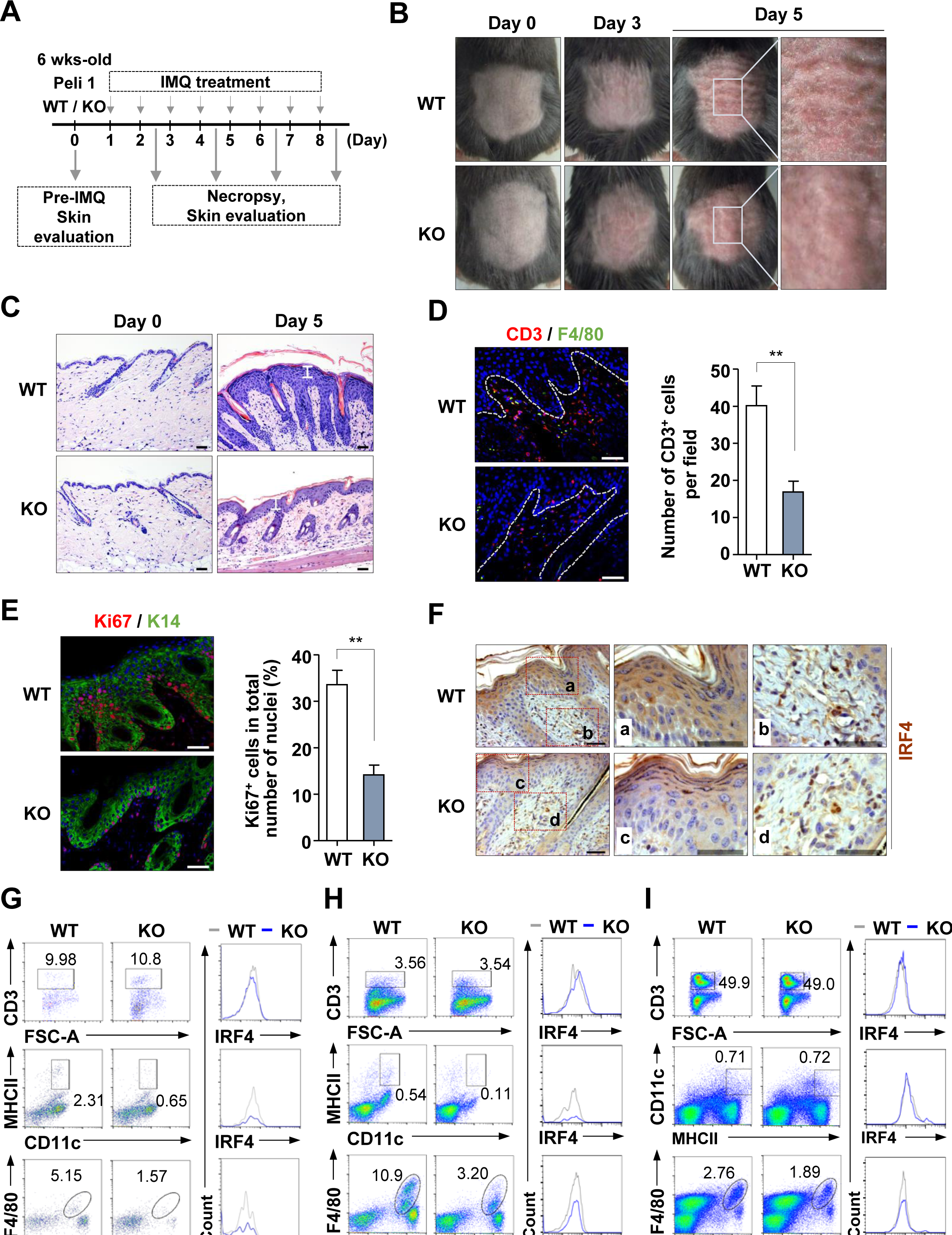
Inhibition of Peli1 delays psoriasis development potentially due to lack of IRF4 induction in psoriatic skin microenvironment. **A.** Timeline showing IMQ trea™ent and necropsy schedules. Peli1 wild type (WT) and Peli1 knockout (KO) mice at six weeks of age were shaved on the back and received daily topical applications of IMQ (62.5 mg). **B.** Psoriasiform lesions in Peli1 WT and KO mice treated with IMQ cream for four days. **C.** Histological presentation of Peli1 WT and KO mice psoriasiform lesions induced by IMQ application at day 5 (n = 5 mice per genotype). White distance bar: 50 μm. **D.** Immunofluorescence staining of T cells (CD3, red) and macrophages (F4/80, green) in lesional skin samples and quantification of CD3^+^ T cells infiltration of Peli1 WT and KO mice (n = 3 mice per genotype). **E.**Immunofluorescence staining to analyze keratinocyte proliferation using keratinocytes (keratin 14, green) and proliferation (Ki67, red) markers. Bar graph shows percentage of Ki67^+^ in K14^+^ cells in skin sections from mice of indicated group (n = 3 mice per genotype). Data are presented as means ± SEM. * *P* < 0.05, ** *P* < 0.01, *** *P* < 0.001, Student’s *t*-test. Scale bars, 50 μm. **F.** Immunostaining of skin sections from IMQ-treated Peli1 WT and KO mice with IRF4 antibody. Scale bars, 50 µm. **G-I.** Representative FACS plots of single-cell suspensions of epidermal (**G**), dermal (**H**), and draining lymph-nodes (**I**) from IMQ-treated Peli1 WT and KO mice. Cells were stained with antibodies specific for CD3, CD11c, MHCII, CD11b, F4/80, and IRF4. Histograms show expression of IRF4 on T cells (CD3^+^, top), DCs (CD11c^+^/MHCII^+^, middle), and macrophages (CD11b^+^/F4/80^+^, bottom). The percentage of each population is shown in plots.

### Peli1 directly binds to IRF4 and causes lysine 63-linked ubiquitination of IRF4

To determine the mechanism involved in the regulation of IRF4 by Peli1, we examined whether there were direct protein-protein interactions between Peli1 and IRF4. Cellular extracts from human primary keratinocyte HaCaT cells were incubated with glutathione S-transferase (GST) or GST-Peli1 fusion protein (Fig 7A). Interaction between GST-Peli1 and endogenous IRF4 was detected, indicating that Peli1 could form a complex with IRF4. Cellular extracts were further immunoprecipitated with an anti-Peli1 antibody or normal IgG followed by immunoblotting with an anti-IRF4 antibody. Consistently, high levels of Peli1 were observed in the complex formed with IRF4 (Fig 7B). A subsequent *in vitro* binding assay using purified His-Peli1 and GST-IRF4 revealed that Peli1 directly interacted with IRF4 (Fig 7C). We then examined whether Peli1-mediated ubiquitination of IRF4 could lead to its stability. Expression plasmid encoding Myc-tagged Peli1 full length (FL), deletion mutant of C-terminal RING-like domain (ΔC), or RING-like domain (C) was transfected into HeLa cells in combination with Flag-IRF4 and HA-tagged ubiquitin WT (HA-Ub) expression plasmids (Fig 7D). Our results showed that overexpression of Peli1 FL clearly induced a high-molecular-mass species of IRF4 which was ubiquitinated (IRF4-Ub). However, overexpression of Peli1 ΔC or Peli1 C did not, indicating that IRF4 ubiquitination depended on Peli1 ubiquitin ligase activity.

**Figure 7.**
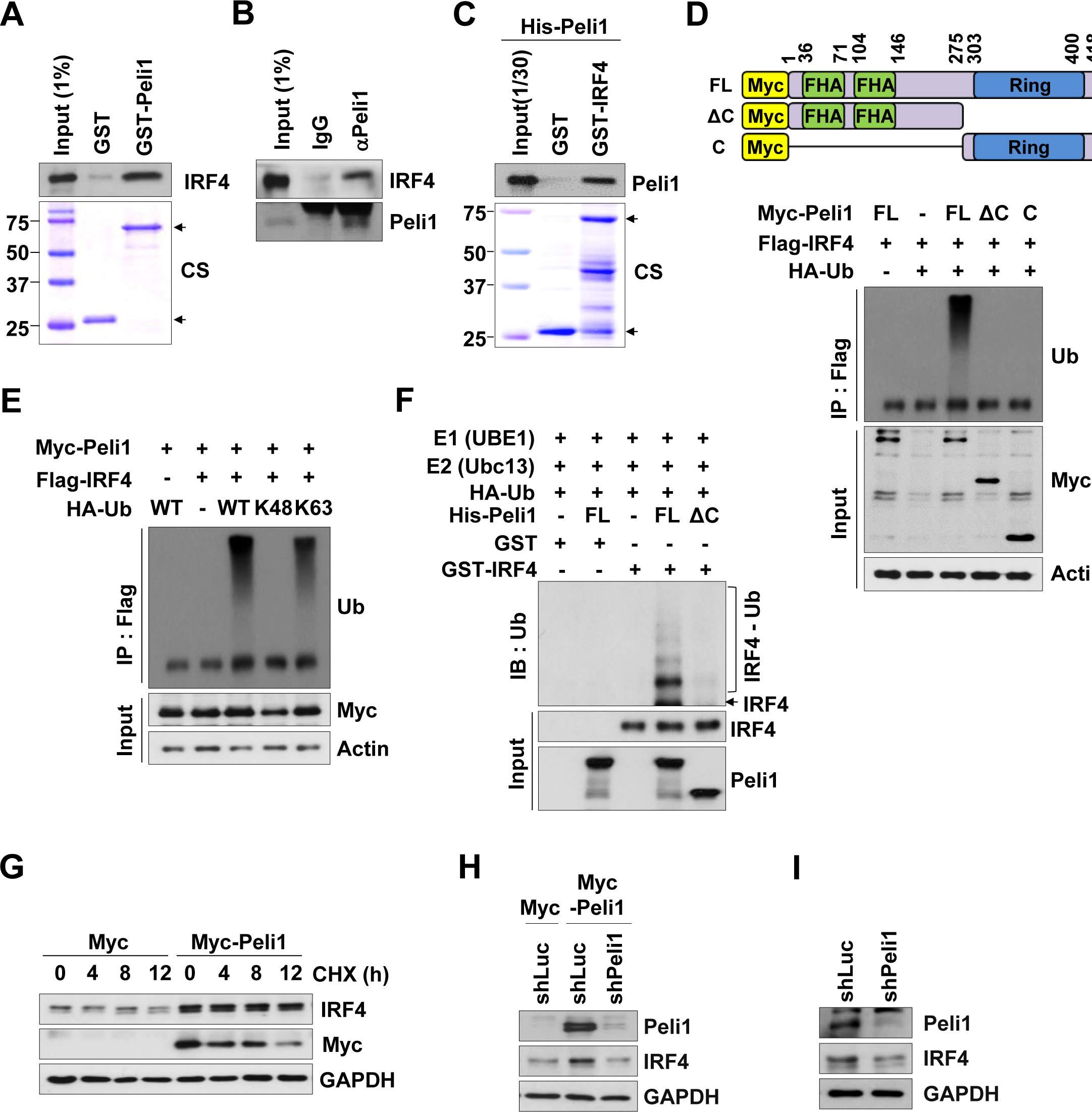
Peli1 interacts with IRF4 and induces lysine 63-linked ubiquitination of IRF4. **A.** GST pull down assay showing IRF4 binding to Peli1 *in vitro*. HaCaT cell lysates were incubated with GST (lower arrow) or GST-Peli1 (upper arrow). Bound proteins were resolved by SDS-PAGE. CS, Coomassie brilliant blue staining. **B.** Immunoblot analysis after co-immunoprecipitation assay using Peli1 antibody. Endogenous Peli1 from HaCaT cell extracts was immunoprecipitated with Peli1 antibody. Co-precipitated endogenous IRF4 was detected with an IRF4 antibody by Western blotting. **C.** *In vitro* binding assay for determination of Peli1 and IRF4 interaction. GST (lower arrow) or GST-IRF4 (upper arrow) was incubated with purified His-Peli1 and subjected to immunoblotting with Peli1 antibody. CS, Coomassie brilliant blue staining. **D.** HeLa cells were transfected with Myc-Peli1 full length (FL), Myc-Peli1 ΔC, or Myc-Peli1 C in combination with HA-Ub and Flag-IRF4 expression plasmids. After transfection, cells were harvested for immunoprecipitation with an anti-Flag antibody. IRF4 protein complex was subjected to immunoblotting with an anti-Ub antibody. A graphical representation of Myc-tagged full-length Peli1 and Peli1 truncation mutants used in *in vitro* and *in vivo* ubiquitination experiments is shown. FHA: FHA domain; RING: RING-like domain. **E.** *In vivo* ubiquitination assay of HeLa cells transfected with Flag-IRF4, Myc-Peli1, and HA-ubiquitin WT, K48, or K63 plasmids as described in methods. Cells were lysed and immunoprecipitated with Flag antibody. Ubiquitinated IRF4 was shown after immunoblotting with anti-Ub. **F.** *In vitro* ubiquitination assay of GST-IRF4 using recombinant E1 (UBE1), recombinant E2 (Ubc13), HA-ubiquitin, and His-Peli1 (FL or ΔC). Reaction mixtures were incubated at 37 °C in assay buffer containing ATP for 2 hours. **G.** Representative cycloheximide chase assay of IFR4 turnover in HaCaT cells transfected with control vector (pMyc) or a vector encoding Myc-tagged Peli1 (pMyc-Peli1). Whole cell lysates were harvested at various time points after exposure to cycloheximide and IRF4 levels were determined by Western blot analysis. **H, I.** HaCaT cells were transfected with control luciferase shRNA (shLuc) or Peli1 shRNA (shPeli1) encoding construct in combination with Myc or Myc-Peli1 expression plasmid. Endogenous IRF4 was detected by Western blotting.

To determine the type of lysine link for the ubiquitination of IRF4 by Peli1, expression plasmid encoding HA-Ub WT, lysine 48 (K48), and lysine 63 (K63) was transfected in combination with Myc-Peli1 and Flag-IRF4 expression plasmids into HeLa cells (Fig 7E). Clearly, Peli1 promoted ubiquitination of IRF4 in the presence of HA-Ub K63. Ubiquitination of IRF4 was then reconstituted *in vitro*. Purified His-tagged Peli1 FL or ΔC protein was incubated with GST or GST-IRF4 in the presence of purified E1, E2, and HA-Ub WT. As shown in Fig 7F, ubiquitinated form of GST-IRF4 became evident after incubation with Peli1 FL, but not with Peli1 ΔC. Taken together, these results demonstrate that Peli1 can directly bind to IRF4 and induce stabilization of IRF4 by K63-linked ubiquitination. To further test whether Peli1 could regulate IRF4 turnover, we transfected HaCaT cells with plasmid encoding Myc-Peli1 in the presence of protein synthesis inhibitor cycloheximide (CHX) (Fig 7G). As expected, overexpression of Peli1 resulted in increased abundance and stability of endogenous IRF4. When we further depleted or overexpressed Peli 1 in HaCaT cells, Peli1 expression directly affected IRF4 stability (Fig 7H, I), indicating that Peli1 could facilitate IRF4 turnover. Our data again strongly suggest that Peli1-directed IRF4 ubiquitination could be a high-risk systemic process in psoriasis microenvironment.

## Discussion

Sequential signaling connections between effectors of innate and adaptive immune systems shape the psoriatic inflammatory process. Activation of Peli1 seems to be a common mediator for instructing psoriatic intercellular signaling pathway from keratinocytes to DCs and subsequently Th17 and Th1 cells (Fig 8). Psoriatic keratinocytes are rich sources of antimicrobial peptides that can trigger the activation of intracellular signaling cascades to DCs and T cells (Buchau & Gallo, 2007; Nestle et al., 2009). Psoriatic keratinocytes activated by Peli1 can stimulate DCs and macrophages to secrete mediators such as IL-21 and IL-23, leading to the differentiation of Th17 and Th1 cells. These T cells in turn secrete mediators (e.g., IL-17a and IL-22) and produce proinflammatory cytokines, chemokines, and others that feed back into the psoriatic inflammatory cycle (Fig 8). By identifying a common intrinsic mechanism in the psoriasis microenvironment that links Peli1-IRF4 signaling axis with the initiation of psoriasis skin inflammation and adaptive immunity, our study provides a novel molecular signaling pathway to better understand the pathogenesis of psoriasis.

**Figure 8.**
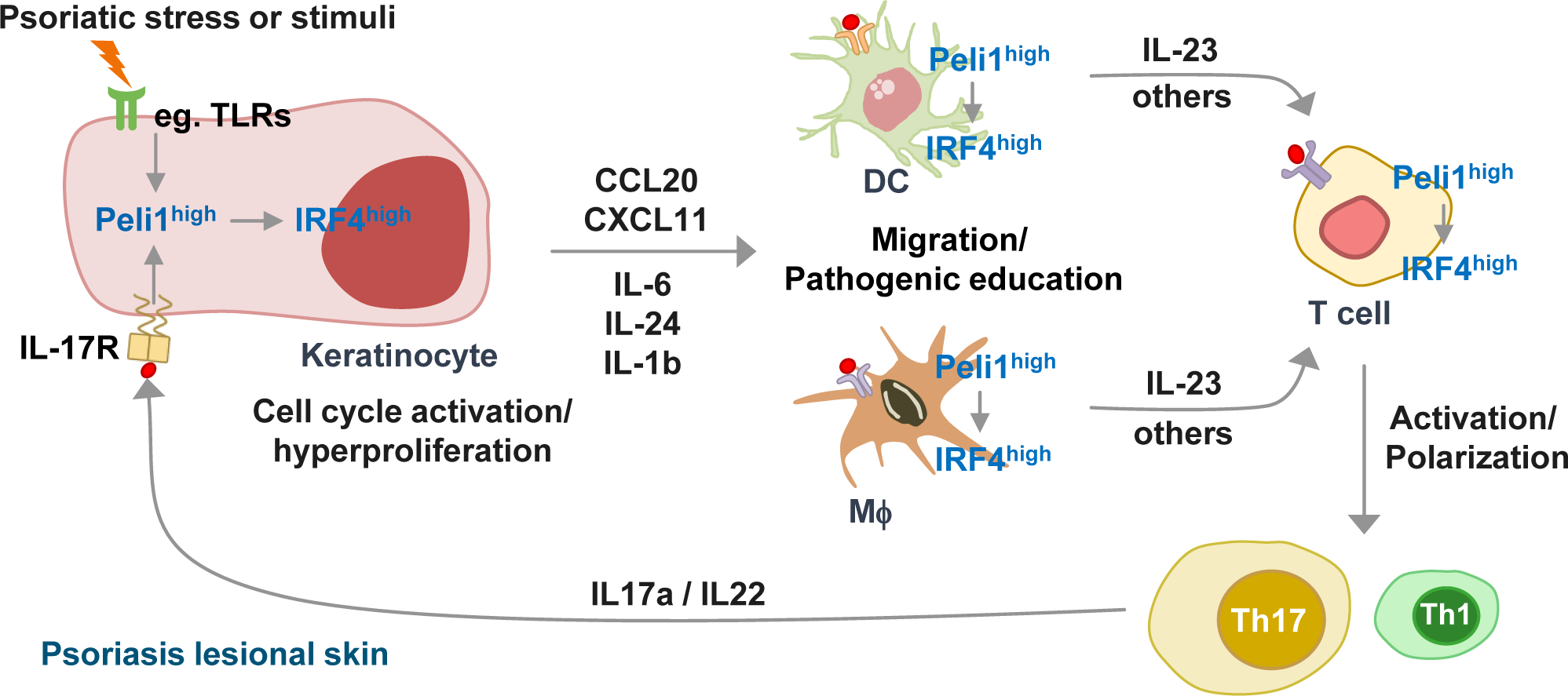
Model of Peli1-mediated IRF4 ubiquitination that generates a feedback cross-link between keratinocyte and T cell response in psoriasis microenvironment. Detailed discussion of this model is provided in the text. DCs: dendritic cells; Mϕ: macrophages.

Receptor-mediated signaling pathways are involved in the regulation of keratinocyte reactions to stress, injury, and infection. Deregulation of surveillance mechanisms in keratinocytes responding to environmental challenges could be a major pathogenic factor triggering the initiation of psoriatic inflammation. Peli1 is a receptor signal responsive E3 ubiquitin ligase that is highly suppressed under normal circumstances. Interestingly, Peli1 expression is differentially regulated by distinct receptor signaling pathways. For instance, stimulation of TCR-CD28-mediated signaling induces Peli1 expression (Chang et al., 2011; Moynagh, 2011). TLR3 and TLR4 signaling can also activate Peli1 expression accompanied by promotion of various signaling cascades such as TRAF6-IRF7 (Humphries & Moynagh, 2015), IKK/TBK1-IRF3 (Smith *et al*, 2011), and IAP2-TRAF3 (Xiao *et al*, 2013). Similarly, IRF4 expression is primarily induced by antigen receptor engagement, stimulation with LPS, or signaling induced by CD40-or IL-4 (Huber & Lohoff, 2014). In T cells, IRF4 is strongly induced within a few hours after TCR stimulation. Its expression is then declined when cells return to a resting state, very similar to patterns of Peli1 expression. TLR and TCR signaling induce Peli1 expression (Chang et al., 2011; Chang et al., 2009; Humphries & Moynagh, 2015; Jin et al., 2012; Moynagh, 2014). Its expression across keratinocytes, macrophages, DCs and T cells is tightly associated with IRF4 induction. Dysregulated receptor-mediated signaling response in a pathogenic microenvironment can lead to aberrant activation of the Peli1-IRF4 signal axis which subsequently initiates chronic skin inflammation clinically identified as psoriasis. Therefore, expression or E3 ligase activity of Peli1 should be turned off after short activation of receptor-mediated signaling in the context of selective signal transduction.

In this study, adaptive transfer experiments revealed that Pepboy.1 chimeric mice receiving Peli1-overexpressing BM cells did not show significant signs of a psoriasis-like phenotype, indicating that overexpression of Peli1 only in T cells seemed to be insufficient to trigger the development of psoriasis. It has been reported that Peli1 mediates rhinovirus-induced expression of CXC-chemokine ligand 8 (CXCL8) chemoattractant in bronchial epithelial cells (Bennett *et al*, 2012). In addition, Peli1 is upregulated in neutrophilic asthma patients (Baines et al., 2011). These results suggest that Peli1-induced chemokine production might affect the development of airway inflammation in asthma and respiratory epithelium after viral infections. Interestingly, results of this study revealed that Peli1-mediated hyperproliferative keratinocytes upregulated the expression and secretion of CCL20 and CXCL11 chemokines known to be indispensable for chemoattraction and inducing immature DC and effector/memory T cells in combination with cytokines. Thus, Peli1-mediated hyperactivation of keratinocytes can initiate pathogenic hyperactivation of T cells and subsequently provide feedback cross-talk between keratinocytes and T cells in the development of skin inflammation and psoriasis lesions. This study further suggests that selective inhibition of T cell activation or polarization alone may be insufficient to cure psoriasis. Thus, inhibition of both keratinocyte hyperactivation and polarized Th17 and Th1 cell responses might be an ideal strategy for targeted psoriasis trea™ent. Recently developed biologic agents can also selectively target the IL-23-Th17 cell axis, including cytokines IL-23, IL-17 and its receptors, and IL-22 for treating psoriasis (Lowes et al., 2013). Although TNFα-and IL-17a-targeting drugs have been recently proven to be effective, a targeting strategy that inhibits Peli1 might be a more powerful way to treat psoriasis. Indeed, our unpublished observation showed that systemic delivery of cyclosporine A or methotrexate (effective and widely used medications for treating psoriasis and other inflammatory diseases) significantly improved their anti-psoriatic effects in Peli1-induced psoriasis mouse model, indicating that Peli1 might be a target for psoriasis.

Research on the pathogenesis of psoriasis has been hindered by the lack of appropriate animal models. Imiquimod trea™ent not only induced phenotypic changes consistent with human psoriasis, but also showed a dependence on the IL-23 and IL-17 axis. However, it is an acute model of inflammation without exhibiting chronic or complex psoriatic-disease phenotype. For example, it does not show comorbidities such as arthritis frequently seen in some human psoriasis patients (Nestle et al., 2009). Another comorbidity of psoriasis includes B cell lymphoma (Davidovici *et al*, 2010). Our recent study strongly indicates that Peli1 expression is upregulated in patients with B cell lymphoma and that such upregulation is associated with poor prognosis (Park et al., 2014). Mice overexpressing Peli1 develop a wide range of lymphoid tumors, particularly B cell lymphoma (Park et al., 2014). In the present study, we found that inducible overexpression of Peli1 for less than 6 months caused severe psoriasis and some comorbidities, but not B cell lymphoma. However, we cannot exclude the possibility that inducible Tg mice might develop B cell lymphoma at later periods as shown in conventional Peli1 Tg mice overexpressing constitutively low levels of Peli1.

IRF family genes can negatively regulate TLR signaling that is central to the activation of innate and adaptive immune systems (Honma *et al*, 2005; Negishi *et al*, 2005). Our present study clearly showed that Peli1 could directly regulate the stability of IRF4 by lysine 63-linked ubiquitination. IRF4 seems to override apoptotic cell death following cell cycle checkpoint activation. It is prone to activating cell proliferation (Huang *et al*, 2012; Shaffer *et al*, 2008). Peli1 can also antagonize the process of apoptosis, thus augmenting cell survival, particularly after prolonged mitotic arrest (Park *et al*, 2017). Indeed, Peli1 accelerates cell cycle progression possibly by IRF4 induction. Moreover, the Peli1-IRF4 signaling axis is critical for activating the hyperproliferation of keratinocytes and the migration of DCs and macrophages specialized in instructing IL-17 responses. A recent study has indicated that human psoriasis trea™ent by phototherapy is associated with IRF4 and IL-17 reduction (Alam et al., 2014). Furthermore, IRF4 has a vital regulatory role during cLN-directed migration of CD11b^+^ dDC which is the most abundant DC subset present in the skin. Such an IRF4-regulated DC migration pathway is likely to be critical for the initiation of adaptive immune responses in skin diseases (such as psoriasis and dermatitis) as well as cutaneous infections(Bajana et al., 2012). Thus, inhibiting the Peli1-IRF4 signaling axis could be an effective therapeutic strategy against psoriasis.

Although this study focused on the Peli1-IRF4-IL-17 signaling axis, we could not exclude the possibility that chronic skin inflammatory responses by acute accumulation of Peli1 might be additionally regulated by other receptor-mediated signaling pathways. Recent evidence from Peli1-deficient mice has shown that Peli1 acts as a critical mediator of TRIF-dependent activation of NF-kB in TLR3 and TLR4 pathways (Chang et al., 2009). Interestingly, activation of NF-kB can induce the expression of IRF4 (Ci *et al*, 2009; Sciammas *et al*, 2006). Thus, Peli1 seems to be a potent regulator for the induction of proinflammatory cytokines. However, we cannot exclude the possibility that Peli1 might catalyze ubiquitination of various target proteins by forming different types of ubiquitin chains in the psoriatic microenvironment.

In summary, we identified a novel pathogenic signaling pathway for Peli1-mediated psoriasis facilitated by feedback cross-talk between keratinocyte hyperactivation and Th17 cell responses. Our results suggest the potential for tailored anti-psoriasis intervention by inhibiting Peli1.

## Materials and Methods

### Animal studies

To generate doxycycline-inducible human Peli1-transgenic mice (rtTA-Peli1), we crossed pTRE TetO Myc-Peli1 transgenic mice with R26-M2rtTA mice (B6.Cg-Gt(ROSA)26Sor^™1^(rtTA*M^2)Jae^/J; The Jackson Laboratory). These animals were maintained under specific pathogen-free conditions. All experiments were performed on mice homozygous for both R26-M2rtTA transgene and human Peli1 transgene. Genotyping was performed with Ready-2X-GO polymerase mixture (HelixAmp^™^) using genomic DNA isolated from tail biopsy samples. To induce expression of human Peli1 transgene, 4-week-old mice were provided drinking water containing 2 mg/ml doxycycline (Sigma) and 5 % sucrose (Sigma) for indicated times. Water was protected from light and exchanged every three days.

Using homologous recombination in C57BL/6NTac embryonic stem cells, a cassette including the coding sequence of *LacZ* gene followed by a promoter-driven *neo* marker surrounded by *frt* sites was introduced between exon 1 and exon 2 of *Peli1* by the European Mouse Mutagenesis Consortium (EUCOMM). In addition, both *neo* and exon 2 of *Peli1* were flanked by *loxP* sites. Details of the allele [*Peli1* “knock out-first-reporter tagged insertion” allele, referred as *Peli1*^*™*1^*a*] and targeting strategy based on EUCOMM are shown in Supplementary Figure S1B. To obtain Peli1 knockout (KO) mice, mice carrying *Peli1*^*™*1*a*^ allele were crossed with B6.FVB-Tg(Zp3-Cre)3Mrt/J mice to excise floxed promoter-driven *neo* selection cassette and obtain Peli1^lacZ/+^ mice. After one backcross to eliminate transgene encoding Cre, Peli1^lacZ/+^ line was intercrossed to generate experimental Peli1^+/+^ (referred to as Peli1 WT) and Peli1^lacZ/lacZ^ (referred to as Peli1 KO) littermates. Mice at 6 weeks of age were shaved on the back and received daily topical application of 62.5 mg imiquimod from a commercially available cream (Aldara; 3M Pharmaceuticals) for 6-8 consecutive days.

All animal experiments were conducted in accordance with guidelines of the Institutional Animal Care and Use Committee (IACUC) of Sungkyunkwan University School of Medicine (SUSM). SUSM is accredited by the Association for Assessment and Accreditation of Laboratory Animal Care International (AAALAC International), abiding by the Institute of Laboratory Animal Resources (ILAR) guidelines.

### Severity scoring of skin inflammation

To score the severity of psoriasiform inflammation, an objective scoring system was developed based on clinical Psoriasis Area and Severity Index (PASI). Erythema and scaling were scored on a scale from 0 to 5: 0, none; 1, slight; 2, moderate; 3, marked; 4, severe; and 5, very severe. Acanthosis was scored on a scale from 0 to 5 based on thickening of skin epidermis: 0, 5 -10 μm (normal); 1, 10 - 20 μm; 2, 20 - 30 μm; 3, 30 - 60 μm; 4, 60 – 90 μm; and 5, > 90 μm. Erythema, scaling, and acanthosis were scored independently. Average score (erythema plus scaling plus thickening) was used to measure the severity of psoriasis (scale: 0-5).

### Plasmid construction, cell culture, transfection, reagents, and antibodies

Full-length cDNA sequence of human Peli1 protein was amplified using oligo-dT primers. Peli1 full length (FL), Peli1 ΔC (1-275), and Peli1 C (275-418) were sub-cloned into Myc-, GST-, and His_6_- tagged plasmids, respectively (Park et al., 2014).Human IRF4 cDNA was amplified from pOTB7-IRF4 plasmid (21C Frontier Human Gene Bank) and sub-cloned into HA-or GST-tagged plasmids. HaCaT cells and HeLa cells were maintained with Dulbecco’s Modified Eagle’s Medium (DMEM) supplemented with 10% FBS (Hyclone). For transient transfection, cells were electroporated using a microporator (Digital Biotechnology) according to the manufacturer’s instructions. Sources of chemicals and reagents were as follows: mouse TLR1-9 agonist kit, InvivoGen; PMA and ionomycin, Sigma; MG132 and cycloheximide, A.G Scientific. All antibodies used in this study are listed in Appendix Table S1.

### Flow cytometry

Lymphocytes were obtained from blood, spleen, and lymph-nodes. Erythrocytes were lysed. Single-cell suspensions were prepared in phosphate buffered saline (PBS) and stained with fluorophore-conjugated antibodies. All antibodies used in this study are listed in Appendix Table S2. For intracellular staining, single-cell suspensions were obtained from skin draining lymph-nodes and cultured for five hours with PMA (50 ng/ml; phorbol 12-myristate 13-acetate) plus ionomycin (500 ng/ml). Brefeldin A (Thermo Fisher Scientific) was added during the final four hours of incubation. After stimulation, cells were washed and fixed with intracellular fixation buffer (Thermo Fisher Scientific) followed by cell permeabilization with permeabilization buffer (Thermo Fisher Scientific). Cells were stained with fluorescently-conjugated cytokine antibodies. Data were obtained using Canto II flow cytometer (BD Biosciences) and analyzed with FlowJo software (FlowJo, LLC).

### TCR stimulation and T cell polarization assay

CD4^+^ T cells were isolated from spleens and lymph nodes using MagniSort^™^ Mouse CD4 T cell Enrichment Kit (Invitrogen). Isolated CD4^+^ T cells were resuspended in RPMI medium supplemented with 10% FBS and then incubated on ice for 30 minutes with anti-CD3 (1 µg/ml) plus anti-CD28 (1 µg/ml). Cells were washed once with 1 ml of cold RPMI medium and stimulated by crosslinking at 37 °C with goat antibody to hamster immunoglobulin (2 µg/ml).

For *in vitro* T cell polarization assay, mouse CD4^+^ T cells were isolated from spleens and lymph-nodes of mice using MagniSort^™^ Mouse CD4 T cell Enrichment Kit (Invitrogen) and cultured for five days on plated-bound anti-CD3 (5 µg/ml) and anti-CD28 (2 µg/ml) under Th1 conditions (IL-12 and anti-IL-4; R&D Systems), Th2 conditions (IL-4 and anti-IFNγ; R&D Systems), or Th17 conditions (TGFβ, IL-6, anti-IL-4 and anti-IFNγ; R&D Systems). Reagents and their concentrations used are shown below: IL-12, 20 ng/ml; IL-4, 40 ng/ml; IL-6, 40 ng/ml; TGFβ, 2 ng/ml; anti-IL-4, 10 µg/ml; and anti-IFNγ, 10 µg/ml.

### Bone marrow transplantation

A schematic illustration of bone marrow chimera experiments is shown in Figs. 4a and 4b. Briefly, male recipient mice were exposed to 10 Gy total-body irradiation. Male donor mice were euthanized under CO_2_ asphyxiation. Femur and tibia bones were then isolated from donor mice. Bone marrow cells were flushed out of the femur and tibia bones into sterile RPMI supplemented with 10% FBS. Following isolation, red blood cells were lysed using RBC lysis buffer (Thermo Fisher Scientific) until cell pellet was void of red color. Next, cell suspension was strained through a 70 µm cell strainer and live cell counts were obtained using a hemocytometer after trypan blue staining. Cells were resuspended in Hanks Balanced Salt Solution (HBSS) at a concentration of 25 x10^6^ cells/ml and stored on ice until transplantation (usually less than one hour). Approximately 4-5 hours following irradiation, 200 µl of isolated bone marrow cells were transplanted into irradiated mice via tail vein injection. Following transplantation, mice were fed normal food and acidified antibiotic water for 14 days for recovery. Recipient mice were bled at two weeks after BMT. Cells were then stained for anti-CD45.1 and anti-CD45.2 antibodies to examine engraf™ent efficiency of BMT.

### Quantitative real-time PCR (qRT-PCR)

We performed qRT-PCR as previously described(Park et al., 2014) using gene-specific primer sets (Appendix Table S3) purchased from QIAGEN. Relative gene expression (in triplicates) was assessed after normalization against expression level of glyceraldehyde 3-phosphate dehydrogenase (GAPDH) as reference gene.

### Patients and Immunohistochemistry

Retrospective patient cohort comprised 156 patients with psoriasis having formalin-fixed paraffin-embedded (FFPE) tissues of their primarily biopsied skin lesion samples at Asan Medical Center from 2010 to 2012. Among them, a total of 18 patients diagnosed with psoriasis by clinical and histological evaluation were consecutively included in this study. Additionally, five normal breast skin tissues obtained from reduction surgery were included for the control group. Histopathological features of each case were reviewed by a dermatopathologist and representative histological findings of psoriasis were confirmed for all included cases.

To evaluate Peli1 expression in human psoriatic lesions, immunohistochemical staining (IHC) was performed with anti-mouse polyclonal Pellino1 antibody using BenchMark XT automated system (Ventana Medical Systems) according to the manufacturer’s protocol. Expression of Peli1 was assessed at each histological site of the skin based on staining intensity and staining sublocalization: (1) histological sites: corneal, spinous and basal layers of epidermis, inflammatory cells, and endothelial cells;(2) staining intensity: 0, no staining; 1+, weak; 2+, moderate; 3+, strong; (3) staining sub-localization: cytoplasmic and nuclear. This study was approved by Asan Medical Center Institutional Review Board (approval number: 2016-0112).

### Histopathological analysis

Paraffin or frozen tissue blocks of mouse skin biopsies were prepared using routine methods and sectioned to obtain consecutive levels. These sections were stained with hematoxylin and eosin. For immunohistochemistry analyses of tissues from mice, sections were deparaffinized, subjected to antigen retrieval in citrate buffer, stained with antibodies, and then incubated with an avidin-biotin-horseradish peroxidase complex (Vectastain Elite ABC kit; Vector Laboratories). Finally, peroxidase activity was visualized using a 3,3’-diaminobenzidine substrate kit (Vector Laboratories). Tissue sections were counterstained with Harris Hematoxylin (BBC Biochemical). For immunofluorescence staining, frozen sections were fixed with 4% PFA, permeabilized with 0.2% tween-20 in PBS, blocked in 5% horse serum, stained with primary antibodies, and incubated with goat anti-rat Alexa 568 and goat anti-rabbit 488 antibodies (Thermo Fisher Scientific). Nuclei were stained with DAPI. Images were taken with a DP72 digital camera mounted onto a BX51 microscope (Olympus Corp).

### Immunoblot, immunoprecipitation, and *in vitro* binding assay

Whole-cell lysates were prepared in NP-40 lysis buffer (150 mM NaCl, 20 mM HEPES, 5 mM EDTA, 0.5% Nonidet P-40, 1 mM PMSF, 1 mM NaF, 1 mM Na_3_Vo_4_, 1 mM DDT) supplemented with a mixture of protease inhibitors and subjected to immunoblot analysis or coimmunoprecipitation assay as described previously(Park et al., 2014). For *in vitro* binding assay, GST fusion proteins were adsorbed onto glutathione-protein A/G-Sepharose beads (Amersham Bioscience) and incubated with HaCaT cell lysates or purified His-Peli1 protein. Bound proteins were separated by sodium dodecyl sulfate polyacrylamide gel electrophoresis (SDS-PAGE) and analyzed by immunoblotting.

### *In vitro* ubiquitination assay

Purified GST or GST-IRF4 (1 μg) was incubated with purified His-Peli1 FL or His-Peli1 ΔC (0.1 μg) in conjunction with E1 (50 ng UBE 1; Boston Biochem), E2 (200 ng UBE2N/Uev1a and 200 ng UbcH5c; Boston Biochem), and HA-tagged ubiquitin (200 ng HA-Ub WT; Boston Biochem) in ubiquitin reaction buffer (5 mM Tris-HCl, 2 mM MgCl_2_, 2 mM ATP, and 100 mM NaCl). Ubiquitin reaction mixtures were incubated at 37 °C for 2 hours followed by immunoblotting. Expression plasmid encoding HA-tagged ubiquitin K48 or K63, in which most lysine residues were mutated to arginine except lysine 48 (K48) or 63 (K63), was kindly provided by Dr. Hong-Tae Kim (Sungkyunkwan University).

### Statistical analysis

Data are presented as means ± SEM. All data were analyzed using GraphPad Prism 4.5 software package (GraphPad Software). A *P* value of less than 0.05 was considered statistically significant. Each experiment was carried out three or more times and reproducible results were obtained. Representative data are shown in figures.

## Acknowledgements

We would like to thank Dr. Seung Woo Lee (POSTECH) for critical reading the manuscript and Drs. Jae Youl Cho (Sungkyunkwan University), Hong-Tae Kim (Sungkyunkwan University), and Ho Chul Kang (Ajou University) for supplying some materials. This study was supported by grants (2011-0030043, 2017R1A2B3006776 & 2017M2A2A7A01070267) of National Research Foundation (NRF) funded by the Ministry of Education, Science, and Technology (MEST), Republic of Korea.

## Author contributions

S. Kim designed and performed the research, prepared figures, and wrote the manuscript. S. Bae, J. Park, and G. H. Ha participated in data generation and analysis.K. Hwang and J. H. Ji provided materials and participated in data generation. H. Go designed studies, analyzed Peli1 expression in psoriasis patient skin samples, and wrote part of manuscript. C. W. Lee designed studies, supervised the overall project, wrote the manuscript, and performed final manuscript preparation. All authors provided feedback and approved the final version of the manuscript.

## The paper explained

### Problem

Chronic skin inflammation including psoriasis is a multisystem disease and is associated with high morbidity and impaired quality of life. In recent years, substantial advances have been made in elucidating molecular mechanisms of psoriasis, resulting in several new treamtents targeting IL-23/T helper cell 17 axis. However, major issues remain to be elucidated, including the primary nature of the disease as an epithelial or immunologic disorder, autoimmune cause of the inflammatory process, the relevance of cutaneous versus systemic factors, and its response to therapy.

### Results

We discovered a novel immunopathogenic intercellular signaling process underlying the connection between innate and adaptive immunity. Our study showed that Peli1 ubiquitin E3 ligase was highly up-regulated in human psoriatic lesions. Our gain-of-function analyses revealed that Peli1 contributed to the development of psoriasis-like disease by stimulating hyperproliferation of keratinocytes and promoting sequential engagement of Th17 cell response. Moreover, Peli1-mediated ubiquitination signaling appeared to be a common systemic intercellular response shared by lesional keratinocytes, dendritic cells, macrophages, and T cells, generating a feedback relationship between keratinocyte and Th17 cell responses in psoriatic microenvironment. Finally, inhibition of Peli1 attenuated immunopathogenic signaling in psoriasis and showed clear signs of recovery from psoriasis.

### Impact

Our study offers Peli1 as a critical mediator for immunopathogenic intercellular signaling in psoriasis microenvironment. Targeting Peli1 could be used as a potential strategy for psoriasis and potentially other chronic inflammatory disease treatments.

